# Rapid contextualization of fragmented scene information in the human visual system

**DOI:** 10.1101/2020.01.06.895870

**Authors:** Daniel Kaiser, Gabriele Inciuraite, Radoslaw M. Cichy

**Affiliations:** Department of Psychology, University of York, York, UK; Department of Education and Psychology, Freie Universität Berlin, Berlin, Germany; Berlin School of Mind and Brain, Humboldt-Universität Berlin, Berlin, Germany; Bernstein Center for Computational Neuroscience Berlin, Berlin, Germany

**Keywords:** visual perception, scene representation, spatial schema, EEG, representational similarity analysis, deep neural networks

## Abstract

Real-world environments are extremely rich in visual information. At any given moment in time, only a fraction of this information is available to the eyes and the brain, rendering naturalistic vision a collection of incomplete snapshots. Previous research suggests that in order to successfully contextualize this fragmented information, the visual system sorts inputs according to spatial schemata, that is knowledge about the typical composition of the visual world. Here, we used a large set of 840 different natural scene fragments to investigate whether this sorting mechanism can operate across the diverse visual environments encountered during real-world vision. We recorded brain activity using electroencephalography (EEG) while participants viewed incomplete scene fragments at fixation. Using representational similarity analysis on the EEG data, we tracked the fragments’ cortical representations across time. We found that the fragments’ typical vertical location within the environment (top or bottom) predicted their cortical representations, indexing a sorting of information according to spatial schemata. The fragments’ cortical representations were most strongly organized by their vertical location at around 200ms after image onset, suggesting rapid perceptual sorting of information according to spatial schemata. In control analyses, we show that this sorting is flexible with respect to visual features: it is neither explained by commonalities between visually similar indoor and outdoor scenes, nor by the feature organization emerging from a deep neural network trained on scene categorization. Demonstrating such a flexible sorting across a wide range of visually diverse scenes suggests a contextualization mechanism suitable for complex and variable real-world environments.

---

The visual world around us is structured in predictable ways at multiple levels (Kaiser et al., 2019a). Natural scenes are characterized by typical distributions of low- and mid-level visual features (Geisler, 2008; Oliva & Torralba, 2003; Purves et al., 2011), as well as typical arrangements of high-level contents across the scene (Bar, 2004; Kaiser et al., 2019a; Torralba et al., 2006; Oliva & Torralba, 2007; Vo et al., 2019; Wolfe et al., 2011). The visual system has adapted to this structure: when multiple scene elements are arranged in a typical way, cortical processing is more efficient (Abassi & Papeo, 2020; Baldassano et al., 2016; Bilalic et al., 2019; Gronau et al., 2008; Kim & Biederman, 2011; Kim et al., 2011; Kaiser et al., 2014, 2020; Kaiser & Peelen, 2018; Roberts & Humphreys, 2010). Such results suggest that when multiple scene elements need to be processed concurrently, cortical processing is strongly tuned to the typical composition of these elements.

In real-life situations, however, we usually do not have access to detailed visual information about all scene elements at once. Instead, visual inputs are fragmented, and only incomplete snapshots of the world are available for visual analysis at any given moment in time. How does the brain assemble a coherent image of the world from such fragmented inputs? To solve this problem, the visual system may draw from internal representations of typical scene structure – scene schemata (Mandler, 1984; Minsky, 1975, Rumelhart, 1980) – in order to contextualize the fragmented inputs with which it is faced. More specifically, schemata may be used to match fragmented visual inputs with their place in the schema: as a result, fragmented visual information should be sorted according to its typical location within the environment. This sorting may help to efficiently contextualize visual inputs.

A recent study showed that incomplete inputs – fragments of natural scenes – are indeed sorted according to their typical location in real-world environments (Kaiser et al., 2019b): In the occipital place area and after 200ms of vision, representations of scene fragments were organized by their typical vertical location in the world. For instance, fragments that typically appear in the upper part of a scene (e.g., a house roof or the ceiling of a room) were represented more similarly to each other than to fragments that typically appear in the lower part of a scene (e.g., a lawn or the room’s floor). No such organization was found for the fragments’ horizontal location, for which clear schemata are missing (Mandler & Parker, 1976).

As a critical limitation, our previous study (Kaiser et al., 2019b) only comprised six different scenes. However, for this mechanism to be useful in the real world, it has to operate across huge amounts of vastly different scenes encountered in our everyday lives. We therefore set out to replicate our findings across a larger and more diverse set of scene images. Here, we used a set of 210 indoor and outdoor scenes, which we split into 4 position-specific fragments each, yielding 840 unique scene fragments (Figure 1a). During an EEG experiment, participants viewed each fragment centrally and in isolation (Figure 1b). Using representational similarity analysis (RSA; Kriegeskorte et al., 2008), we then tracked the fragments’ cortical representations across time. As the key result, we found that within the first 100ms of visual processing and most prominently after 200ms, the fragments’ cortical representations were organized by their vertical location within the full scene. This neural organization was neither explained by visual similarities amongst indoor and outdoor scenes, nor by the features extracted by a deep neural network (DNN) trained on scene categorization, suggesting that it is not immediately explicable by differences in simple visual features. We conclude that the visual system uses scene schemata to sort inputs according to their typical location within the environment, supporting the contextualization of fragmented visual information.

**Figure 1.**
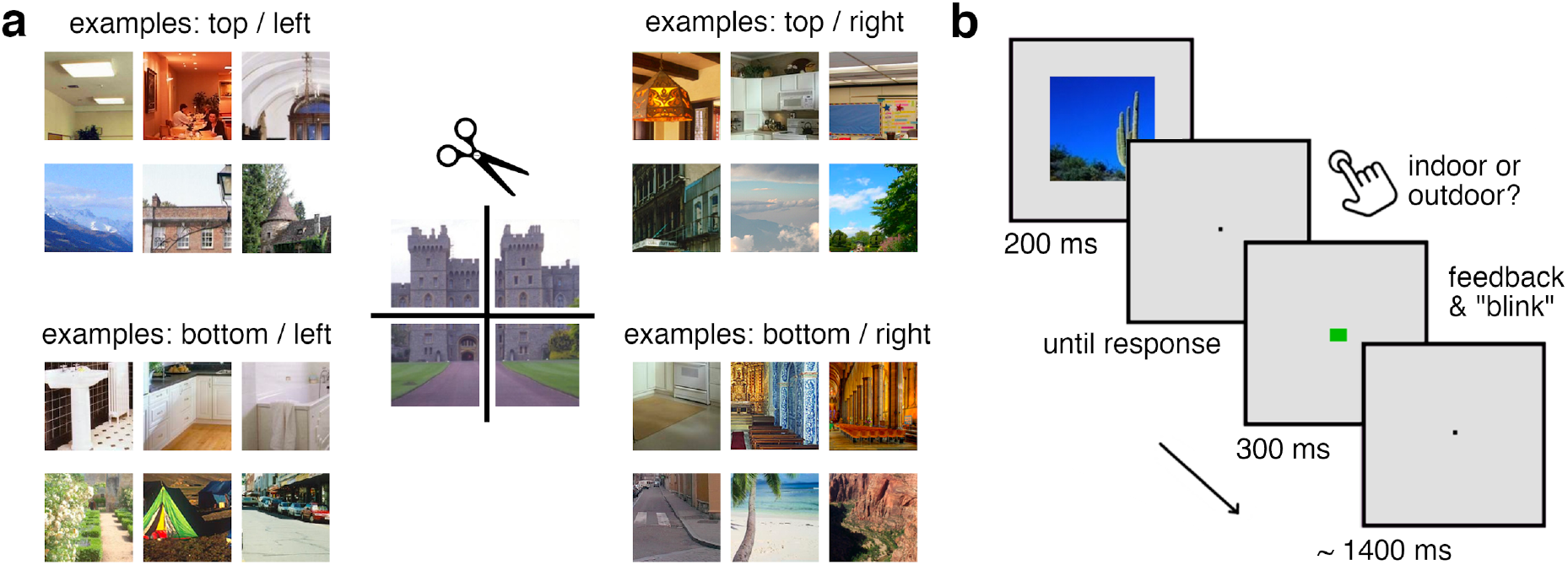
Stimuli and paradigm. a) To mimic the fragmented nature of natural visual inputs, we used a set of 210 widely varying indoor and outdoor scene photographs, from a total of 14 categories. Each scene was split into four equally sized fragments (top/left, top/right, bottom/left, bottom/right). The panel shows representative examples of different indoor (upper rows) and outdoor (lower rows) scenes. b) During the EEG experiment, participants viewed the individual fragments in the center of the screen while performing an indoor/outdoor categorization task.

## Materials and Methods

### Participants

Twenty healthy adults (mean age 27.3, *SD*=4.6; 12 female) participated in the study. The sample size was identical to the sample size of our previous EEG study (Kaiser et al., 2019b). All participants had normal or corrected-to-normal vision. Participants provided informed consent and received monetary reimbursement or course credits. All procedures were approved by the ethical committee of Freie Universität Berlin and were in accordance with the Declaration of Helsinki.

### Stimuli

Stimuli were 210 natural scene photographs, taken from an online resource (Konkle et al., 2010). Half of the stimuli depicted outdoor scenes from seven different categories (bridges, camping sites, historical buildings, houses, nature scenes, streets, and waterfronts) and half of the stimuli depicted indoor scenes from seven different categories (bathrooms, bedrooms, churches, classrooms, dining rooms, kitchens, and living rooms). To create position-specific fragments, each scene was split along the vertical and horizontal axes (Figure 1a), yielding four fragments of equal size for each scene and 840 fragments in total. The full stimulus set is available on OSF (doi.org/10.17605/OSF.IO/D7P8G). During the experiment, these fragments were presented individually and in the center of the screen (5.5° by 5.5° visual angle). Participants were not shown the full scene images prior to the experiment.

### Experimental paradigm

During the experiment, participants briefly viewed the individual scene fragments, all presented in the same central location (Figure 1b). Each of the 840 fragments was shown twice during the experiment, yielding 1,680 trials. Trial order was randomized separately for the first and second half of trials, so that every fragment appeared once in the first half of the experiment and once in the second half. On each trial, a single fragment appeared for 200ms and participants were asked to categorize the fragment as either stemming from an indoor scene or an outdoor scene using two keyboard buttons. After every response, the fixation cross turned red or green for 300ms, indicating response correctness. Trials were separated by an inter-trial interval varying randomly between 1,300ms and 1,500ms. Participants performed the categorization task well (93% correct responses, *SE*=1%; 769ms average response time, *SE*=36ms), with no differences in accuracy or response time between fragments stemming from the top versus the bottom or from the left versus the right (all *t*[19]<1.89, *p*>0.07). Further, participants were instructed to maintain central fixation throughout the experiment, and to only blink after they had given a response. Stimulus presentation was controlled using the Psychtoolbox (Brainard, 1997).

### EEG recording and preprocessing

EEG signals were recorded using an EASYCAP 64-electrode system and a Brainvision actiCHamp amplifier. Electrodes were arranged in accordance with the 10-10 system. EEG data were recorded at 1000Hz sampling rate and filtered online between 0.03Hz and 100Hz. All electrodes were referenced online to the Fz electrode. Offline preprocessing was performed using FieldTrip (Oostenveld et al., 2011). EEG data were epoched from −200ms to 800ms relative to stimulus onset, and baseline-corrected by subtracting the mean pre-stimulus signal. Channels and trials containing excessive noise were removed based on visual inspection; on average, 1.5 channels (*SD*=0.6) and 159 trials (*SD*=84) per participant were removed. Blinks and eye movement artifacts were removed using independent component analysis and visual inspection of the resulting components. The epoched data were downsampled to 200Hz.

### Measuring representational similarity

To track the representations of individual fragments across time, we used representational similarity analysis (RSA; Kriegeskorte et al., 2008). First, we created neural representational dissimilarity matrices (RDMs) for each time point in the EEG epochs (5ms resolution), reflecting the pairwise dissimilarity of the fragments’ brain representations. Second, we modeled the organization of the neural RDMs in a regression approach (Kaiser et al., 2019b; Proklova et al., 2016, 2019), which allowed us to track when representations are explained by the fragments’ vertical and horizontal location within the full scene as well as the scene’s category.

To construct neural RDMs, we computed the pairwise dissimilarity of all fragments at each time point using the CoSMoMVPA toolbox (Oosterhof et al., 2016). This analysis was done separately for each participant. For this, we used response patterns across 17 posterior electrodes (Kaiser et al., 2019b) in our EEG montage (O1, O2, Oz, PO3, PO4, PO7, PO8, POz, P1, P2, P3, P4, P5, P6, P7, P8, Pz); results for central and anterior electrode groups can be found in Supplementary Figure S1. For each of the 840 fragments, we computed the response pattern to the fragment by averaging across the two repetitions of the fragment. If after trial rejection during preprocessing only one trial was left for a fragment, we used the response pattern from this one trial. If after preprocessing no trial was left for a fragment, this fragment was excluded from the analysis (i.e., removed from all RDMs of the respective participant). Neural dissimilarity was computed by correlating the response patterns to each individual fragment in a pairwise fashion and subtracting the resulting correlations from 1, yielding an index of neural dissimilarity (0: minimum dissimilarity, 2: maximum dissimilarity). Computing this index for each pairwise comparison of fragments, we obtained an 840-by-840 neural RDM for each time point.

### Modelling representational similarity

To quantify how well the neural organization is explained by the fragments’ vertical and horizontal location within the full scene and by the original scene’s category, we modeled the neural RDMs in general linear models (GLMs) with three predictors: (1) a vertical location RDM, in which each pair of conditions is assigned either a value of 0, if the fragments stem from the same vertical location (e.g., both from the top), or the value 1, if they stem from different vertical locations (e.g., one from the top and one from the bottom), (2) a horizontal location RDM, in which each pair of conditions is assigned either a value of 0, if the fragments stem from the same horizontal location (e.g., both from the left), or the value 1, if they stem from different horizontal locations (e.g., one from the left and one from the right), and (3) a category RDM, in which each pair of conditions is assigned either a value of 0, if the fragments stem from the same scene category (e.g., both fragments stemming from scenes showing bridges), or the value 1, if they stem from different categories (e.g., one stemming from a bridge and one stemming from a living room).

GLMs were constructed with the neural RDMs as the regression criterion and the vertical and horizontal location RDMs as well as the category RDM as predictors. For these GLMs, the neural RDMs and predictor RDMs were vectorized by selecting all lower off-diagonal elements – the rest of the entries, including the diagonal, were discarded. Values for the neural RDMs were z-scored. Estimating these GLMs yielded three beta weights for each time point and participant. We subsequently tested these beta weights across participants against zero, which revealed whether the fragments’ vertical location, horizontal location, and their category significantly explained the neural organization at each time point.

### Controlling for shared properties among indoor and outdoor scenes

We additionally performed two control analyses. In the first control analysis, we aimed at eliminating visual and conceptual features that are common to either indoor or outdoor scenes (e.g., the top fragments from outdoor scenes often show blue skies). We thus constructed RDMs for horizontal and vertical location information which only contained comparisons across indoor and outdoor scenes. These RDMs were constructed in the same way as explained above, but now all comparisons within the same scene type (e.g., comparisons of different indoor scene fragments) were removed. We then repeated the original GLM analysis (see above) with these restricted RDMs, allowing us to see if the organization according to vertical location persists when only comparisons between indoor and outdoor scenes are considered.

### Controlling for categorization-related visual features

In the second control analysis, we investigated whether visual features related to scene categorization predicted the fragments’ location-specific organization. To model the extraction of visual features during scene processing, we used a deep neural network (DNN). DNNs provide the current state of the art in modelling representations in the human visual system (Cichy & Kaiser, 2019; Kietzmann et al., 2018; Kriegeskorte, 2014), and they capture a variety of features extracted during cortical scene processing (Cichy et al., 2017; Groen et al., 2018). Here, we used a ResNet50 network (He et al., 2016), pretrained on scene categorization using the Places365 dataset (Zhou et al., 2016), as implemented in PyTorch. ResNet50 consists of 4 residual layer modules, each composed of multiple blocks of convolutional layers. These modules are followed by a single fully-connected layer (see Figure 4a for a schematic of the architecture). We ran all fragments through the DNN and extracted RDMs along the network. The entries in these RDMs reflected pairwise distances (1-correlation) between the fragments’ DNN activation vectors obtained from a given layer. In this way, a separate RDM was computed for the final layer of each residual module (layers res2c, res3d, res4f, and res5c) and for the fully-connected layer, yielding 5 RDMs in total.

**Figure 2.**
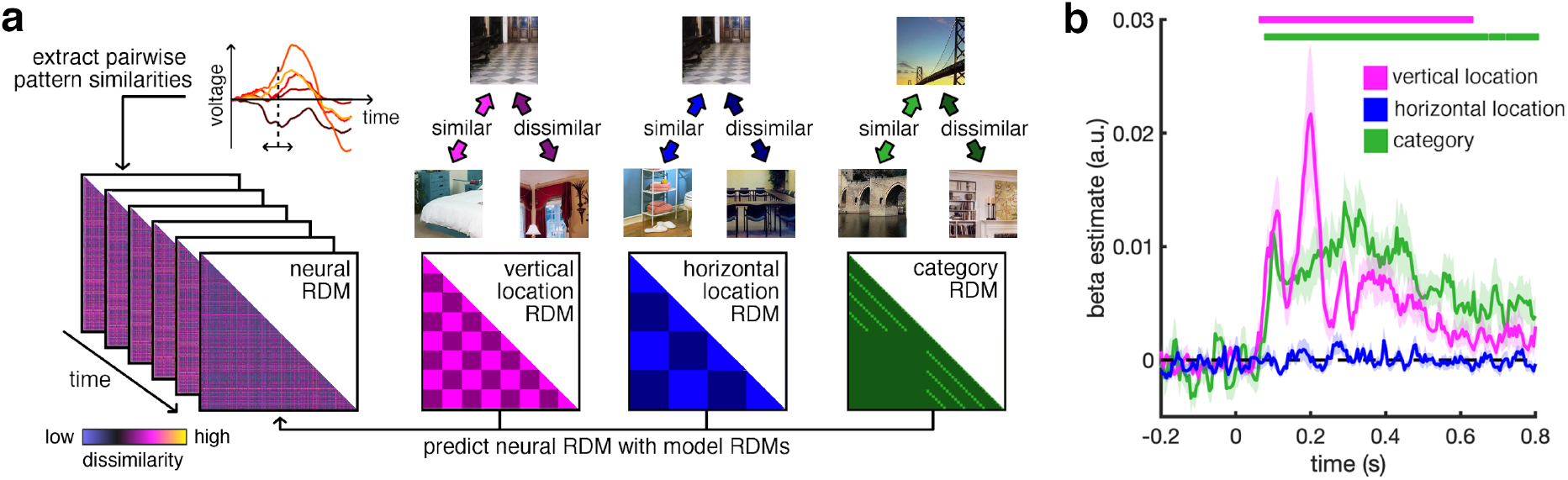
Analysis approach and main result. a) We first extracted neural RDMs from EEG signals in a time-resolved manner. That is, for each time point in an epoch we correlated the response patterns evoked by each one fragment with the response pattern evoked by each other fragment, yielding an 840-by-840 matrix of pairwise neural dissimilarities. The neural RDMs were then modeled as a combination of three predictor RDMs that captured the fragments’ dissimilarity in vertical location (e.g. different fragments from the same vertical location were considered similar), horizontal location (e.g. different fragments from the same horizontal location were considered similar), and category (e.g. different fragments from the same category, such as both from bridges, were considered similar). Estimating this model for each time point yielded three time courses of beta estimates, indicating how well the neural organization matched each of the predicted organizations. b) The fragments’ vertical location (but not their horizontal location) predicted neural organization between 70ms and 625ms, suggesting a sorting of information according to typical real-world locations. Additionally, the fragments’ category predicted their neural organization between 85ms and 800ms. Significance markers denote *p_corr_*<0.05. Shaded margins represent standard errors of the mean.

**Figure 3.**
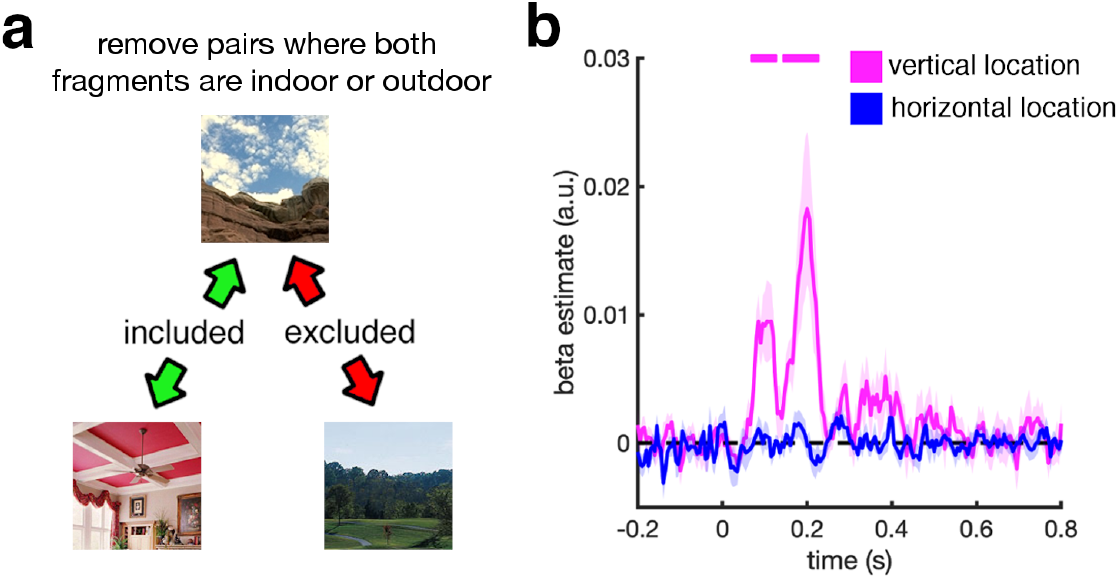
Controlling for visual similarity among indoor or outdoor scenes. a) In this analysis, we removed all pairwise comparisons between the fragments of the same type (i.e., both indoor or both outdoor) from the neural and predictor RDMs. This allowed us to control for visual and conceptual features shared by fragments stemming from the same location (e.g., fragments from the upper part of outdoor scenes often contain skies and clouds). d) Removing these comparisons did not abolish vertical location information, which remained significant between 75ms and 120ms, and between 150ms and 220ms. This indicates that the sorting of fragments according to their vertical location in the world is flexible with regards to visual and conceptual attributes shared among the indoor or outdoor scenes. Significance markers denote *p_corr_*<0.05. Shaded margins represent standard errors of the mean.

**Figure 4.**
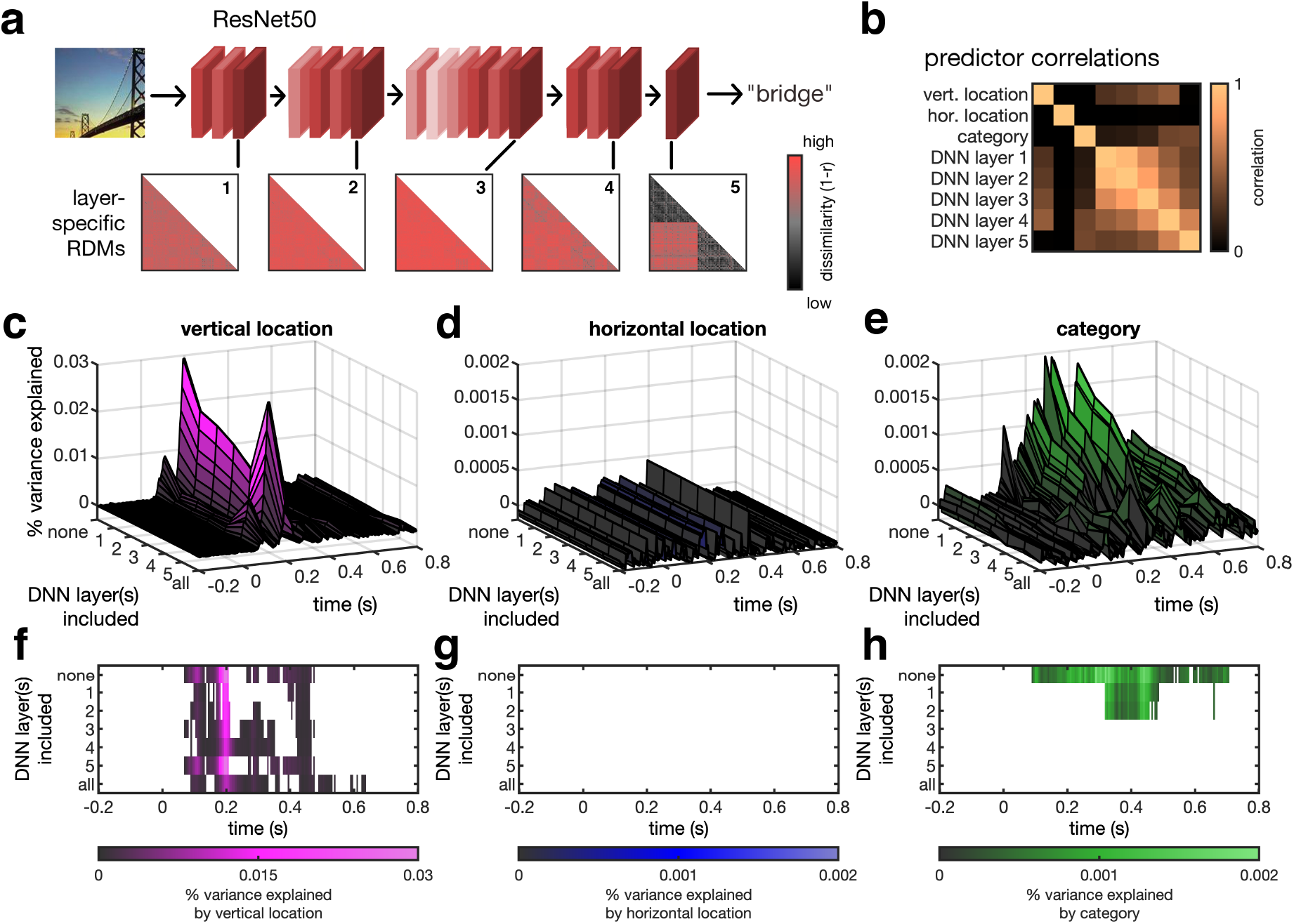
Controlling for categorization-related visual features. a) To quantify the visual features emerging during categorization, we used a ResNet50 DNN pretrained on scene categorization (the panel shows a simplified depiction of the architecture). We ran all 840 fragments during this DNN and extracted RDMs for five different layers at different depths. b) When correlating the different predictor matrices, we found moderate correlations between the DNN RDMs and the vertical location and category predictor RDMs, as well as between the different DNN layers. To establish whether the vertical location and category organizations in the neural data were accurately captured by visual features extracted by the DNN, we performed model comparison analyses. For each of the three predictors (vertical location, horizontal location, and category) separately, we devised two models: a full model, which included all three predictors, and a reduced model, which did not contain the predictor of interest; additionally, both models could contain one of the DNN layers or all the DNN layers. By subtracting the R^2^ value of the reduced model from the R^2^ value of the full model, we obtained the amount of additional variance explained by the predictor of interest. c-e) Additional variance explained by including the vertical location (c), horizontal location (d), and category (e) predictors into the model, as a function of which DNN layers were included in the model. f-h) Thresholded maps of the data in (c-e), where only time points and models are shown where including the predictor adds significant explained variance (*p_corr_*<0.05, corrected for multiple comparisons across time, separately for each model). The model comparison analyses revealed two key insights: First, the fragments’ vertical location predicted additional variance beyond the visual feature organization emerging from different DNN layers, with the same peak structure as in the previous analyses (see Supplementary Table S1 for details). Second, no additional variance was explained by the category predictor when RDMs from deeper layers of the DNN were included as predictors. This shows that although the DNN is a good model for categorization-related visual features, it is unable to fully account for the vertical-location organization in the neural data.

To assess whether RDMs across different depths of the DNN could explain the category and location organizations in the neural RDMs, we performed model comparison analyses. We conducted analyses separately for each participant and for each time point of the epoched EEG data. For each of these analyses, we devised a full model that modelled the neural RDMs as a combination of a set of predictors: (1) the vertical location RDM, (2) the horizontal location RDM, and (3) the category RDM. Additionally, the model could either contain one of the DNN RDMs or all of the DNN RDMs, yielding seven different types of models (no DNN layer included, one of the five DNN layers included, or all DNN layers included). We compared each full model to a reduced model, where the predictor of interest was not included: for instance, to see whether the category RDM added information beyond the vertical location, horizontal location, and DNN RDM(s), we devised a model that contained all predictors from the full model, but not the category predictor. To compare the full model and the reduced model, we computed adjusted R^2^ values (which reflect the amount of variance explained by the model, adjusted for the number of predictors) for both models. By subtracting the R^2^ value of the reduced model from the R^2^ value of the full model, we obtained an index of how much additional variance is explained by the predictor left out in the reduced model.

In Supplementary Figure S2, we additionally report R^2^ values indexing the fit of each of the three key predictors (vertical location, horizontal location, and category) to the data and the fit of the different DNN layers to the data. To put these R^2^ values into perspective, we computed an empirical noise ceiling which reflected the lower bound of explicable variance in the neural data. For every participant, we correlated this participant’s RDM with the average RDM of all other participants. These correlation values were then squared to obtain an R^2^ value for each participant and time point. Averaging across participants yielded a time-resolved empirical noise ceiling.

### Statistical testing

To test whether GLM beta weights (or differences in R^2^ values) were significantly greater than zero, we used a threshold-free cluster enhancement procedure (Smith and Nichols, 2009) and multiple-comparison correction based on a sign-permutation test (with null distributions created from 10,000 bootstrapping iterations), as implemented in CoSMoMVPA (Oosterhof et al., 2016). The resulting statistical maps were thresholded at *z*>1.64 (i.e., *p_corr_*<.05). For all peaks in the time series, we additionally report results of conventional one-sided t-tests against zero. To estimate the robustness of peak latencies we performed a bootstrapping analysis. In this analysis, we created 1000 samples of 20 randomly chosen datasets each (with possible repetitions). For each random sample, we computed the peak latency (i.e., the highest beta estimate in the average time course). We then computed a confidence interval *(ci)* by selecting the central 95% of the distribution across the 1000 random samples. Given the clear two-peak structure in vertical location information, we performed the bootstrapping analysis separately for the early and late peaks, by splitting the data for each random sample along the minimum beta value between 100ms and 200ms.

### Data Availability

Data and stimuli are publicly available on OSF (doi.org/10.17605/OSF.IO/D7P8G). Other materials are available upon request.

## Results

To model the fragments’ cortical representations across time, we ran a GLM analysis with three predictors capturing the fragments’ vertical and horizontal locations within the full scene and the full scene’s category (Figure 2a). This analysis revealed three key insights. First, the fragments’ cortical organization was explained by their vertical location within the scene (Figure 2b), from 70ms to 625ms (peaking at 110ms, peak *t*[19]=4.11, *p*<0.001, *p_corr_*<0.05, *ci*=[85ms, 115ms], and at 200ms, peak *t*[19]=3.54, *p*=0.001, *p_corr_*<0.05, *ci*=[190ms, 205ms]). This suggests that fragmented scene information is sorted by its typical origin within the visual world. Second, the fragments’ horizontal location did not significantly predict their neural organization, suggesting that the more rigid real-world location along the vertical axis is more strongly reflected in cortical signals. Third, the fragments’ category (i.e., which of the 14 scene categories a fragment stemmed from) was also reflected in their neural organization, from 85ms to 800ms (peaking at 290ms, peak *t*[19]=4.94, *p*<0.001, *p_corr_*<0.05, *ci*=[100ms, 447.5ms]). This finding supports previous studies showing that scene category can be rapidly decoded from EEG signals (Dima et al., 2018; Kaiser et al., 2019b; Lowe et al., 2019).

Together, these results suggest that fragmented visual information is organized with respect to its typical vertical location in the world. To test how flexible this organization is with respect to visual and conceptual properties of individual scenes, we additionally performed two control analyses.

In the first control analysis, we tested whether the sorting of fragmented information is independent of visual information shared among the indoor or outdoor scenes. In this analysis, we restricted our models to comparisons between indoor and outdoor scenes (i.e., all comparisons within the same scene type were removed from all RDMs). The comparisons between indoor and outdoor scenes share fewer visual and conceptual properties than the comparisons of scenes from the same type (Figure 3a): For example, two fragments from the upper part of outdoor scenes often share both visual properties (e.g., they tend to be blue-colored) and conceptual content (e.g., they tend to contain the same objects, such as clouds or tree tops).

This analysis revealed significant vertical location information (Figure 3b), from 75ms to 120ms (peaking at 105ms, peak *t*[19]=2.95, *p*=0.004, *p_corr_*<0.05, *ci*=[80ms, 120ms]) and from 150ms to 220ms (peaking at 200ms, peak *t*[19]=3.07, *p*=0.003, *p_corr_*<0.05, *ci*=[190ms, 207.5ms]), but no horizontal location information. This suggests that the cortical sorting of information according to the fragments’ vertical location occurs similarly for visually and conceptually diverse scenes. Note that – as within-category comparisons were removed – no category information could be computed in this analysis.

In the second control analysis, we used a ResNet50 DNN (He et al., 2016) trained on scene categorization to quantify the organization of categorization-related visual features across the fragments. We first computed RDMs for five layers at different depths of the DNN (Figure 4a). These RDMs showed moderate correlations with the vertical location predictor RDM (from layer 1 to 5: r=0.25, r=0.30, r=0.37, r=0.46, r=0.01) and with the category predictor RDM (from layer 1 to 5: r=0.12, r=0.14, r=0.18, r=0.35, r=0.36), but not with the horizontal location predictor RDM (all r<0.02) (Figure 4b). We then performed model comparison analyses, in which we investigated whether the fragments’ vertical location, their horizontal location, and their category explained variance in the neural data, beyond the variance explained by the features extracted by the DNN. In these analyses, we compared full models that contained the DNN predictors and the three key predictors (vertical location, horizontal location, and category) with reduced models that lacked the predictor of interest (e.g., vertical location). Comparing the variance explained by the full model and the reduced model allowed us to quantify how much variance the left-out predictor of interest explained beyond the variance already explained by the other predictors (see Materials and Methods for further details on these model comparison analyses).

When comparing models that contained the vertical location predictor and models that did not contain the vertical location predictor, we found that the inclusion of the predictor improved the model fit for all models tested (Figure 4c): Independently of which DNN layers were included in the model, the difference in the R^2^ values of the full and reduced models remained significant (Figure 4f), with two consistent peaks in vertical location at around 110ms and 200ms (timings and confidence intervals for these peaks can be found in Supplementary Table S1). Critically, even when all DNN layers were included in the model, vertical location explained additional variance between 90ms and 135ms, and again from 165ms. By contrast, horizontal location never explained any additional variance, for all the models tested (Figure 4d/g). Finally, category information did not explain much variance beyond the DNN model (Figure 4e): When layers 1 or 2 were included in the model, including the category predictor explained additional variance between 320ms and 485ms (Figure 4h). However, when deeper layers or all layers together were included in the model, the category predictor added no additional variance beyond the DNN, showing that the DNN accurately captures the visual features that the brain uses for categorization. Remarkably, these features were unable to explain the fragments’ cortical organization according to vertical location.

Together, our results suggest that fragmented information is sorted according to its typical vertical location in the world, providing a mechanism for the contextualization of incomplete visual information. Even when controlling for visual and conceptual scene attributes, this mechanism can rapidly structure the cortical organization of incoming information.

## Discussion

During natural vision the brain is constantly faced with incomplete snapshots of the world from which it needs to infer the structure of the whole scene. Here, we show that in order to meet this challenge, the visual system rapidly contextualizes incoming information according to its typical place in the world: within the initial 100ms of processing and most strongly after 200ms, fragmented scene information is sorted according to its real-world location. By using a large stimulus set (comprising 840 unique fragments) we provide compelling evidence that this mechanism supports spatial contextualization across diverse visual environments.

Which features allow the visual system to make such inferences about a fragment’s typical position within the environment? Across all analyses, the strongest vertical-location organization was found after 200ms of processing. At this time, the amount of variance explained by the vertical location predictor was also highest (see Supplementary Figure S2). Further, this timing solidifies our previous results (Kaiser et al., 2019b), where the strongest vertical location organization in EEG signals became apparent at around 200ms after onset. At this time higher-level scene attributes – such as the scene’s clutter or openness (Cichy et al., 2017; Harel et al., 2016) – are analyzed, suggesting that the sorting of information according to real-world locations is determined by more complex scene properties, rather than low-level visual features. This is consistent with previous fMRI results which demonstrated a vertical location organization in the occipital place area, but not in early visual cortex (Kaiser et al., 2019b).

We additionally found a very rapid onset of vertical location information with the first peak after 100ms. Further, we found that the fragments’ vertical location correlated with visual attributes extracted at different levels of a DNN model, including representations emerging in early processing stages (see Figure 4b). However, contrary to our previous study (Kaiser et al., 2019b), this early effect was not fully explained by categorization-related features as quantified by a DNN model (see Figure 4c/f). This discrepancy may be related to the limited stimulus set in our previous study, which was unlikely to cover all low- and mid-level feature differences that are diagnostic of a fragment’s vertical location in the world. What the current result suggests is that there are visual features that are analyzed early on and which are used to spatially contextualize visual information. Their cortical organization is not fully explained by our DNN model, which suggests that these features are not analyzed in the same way during categorization – instead, they may be particularly useful for spatially contextualizing inputs. Such features could comprise particular distributions of spatial frequency content or texture information (Dima et al., 2018; Groen et al., 2013, 2017). Alternatively, these early effects could reflect the rapid analysis of scene geometry (Henriksson et al., 2019). Future studies need to explicitly isolate the contribution of different visual features to the sorting of visual information by real-world location at different processing times.

Previous research has demonstrated that the representation of individual naturalistic stimuli depends on whether their current position in the visual field matches their typical position in the world (Chan et al., 2010; de Haas et al., 2016; Mannion, 2015; Kaiser & Cichy, 2018; Kaiser et al., 2018). For instance, when face parts (e.g., an eye) or objects (e.g., a lamp) are presented in their typically experienced visual-field position (e.g., the upper visual field), they evoke more efficient cortical representations (de Haas et al., 2016; Kaiser & Cichy, 2018). This suggests that across diverse visual contents cortical representations of a stimulus are entwined with preferences for its typical location (Kaiser & Haselhuhn, 2017; Kaiser et al., 2019a). Here we show that the pairing of visual representations and location information is apparent even when objects do not appear in their expected locations: although in the current study all fragments were presented in the same central location, their representations were still organized by their typical location. This shows that even in the absence of location information the brain can use the characteristic spatial distribution of visual contents to organize their representation in an efficient way. A related effect of real-world structure on individual object representations was previously reported in face processing, where representations of individual face fragments are grouped according to their position within a face (Henriksson et al., 2015). Together, these results suggest that information across different types of fragmented visual contents can be contextualized on the basis of real-world structure.

How does this contextualization mechanism aid perception under naturalistic conditions? The mechanism may be particularly beneficial across a variety of situations where visual inputs are incomplete. Such situations include partially occluded objects, fast-changing and dynamic environments, and fragmented information arising from eye movements across a scene. In each of these situations, matching the input with its typical position in the context of the current environment can facilitate the understanding of the incomplete information available at every point in time. Future studies need to connect the rapid sorting process described here and behavioral benefits in the aforementioned situations.

## Acknowledgements

D.K. and R.M.C. are supported by Deutsche Forschungsgemeinschaft (DFG) grants (KA4683/2-1, CI241/1-1, CI241/3-1). R.M.C. is supported by a European Research Council Starting Grant (ERC-2018-StG 803370). Thanks for Kshitij Dwivedi for help with the DNN model computations. The authors declare no conflict of interest.

## Author Contributions

D. K. and R.M.C. designed research, D.K. and G.I. acquired data, D.K. and G.I. analyzed data, D.K., G.I., and R.M.C. interpreted results, D.K. prepared figures, D.K. drafted manuscript, D.K., G.I., and R.M.C. edited and revised manuscript. All authors approved the final version of the manuscript.

