## Supplementary Information for "Rapid contextualization of fragmented scene information in the human visual system"

Supplementary Contents:


- *Figure S1.* Results for different electrode groups
- *Figure S2.* Explained variance for different representational models
- *Table S1.* Peak timings for vertical location information in the model comparison analyses

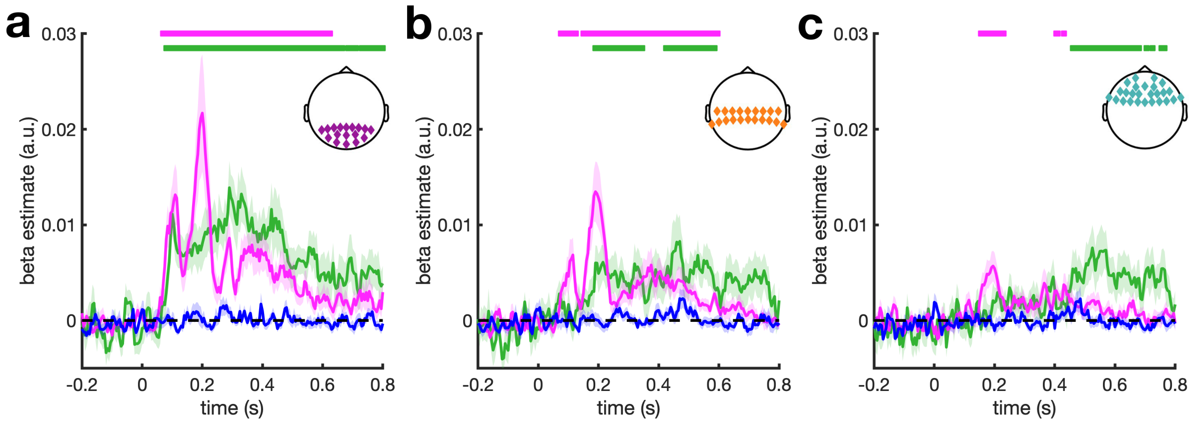


*Figure S1.* Results for different electrode groups. a) In the posterior electrode group, the fragments' vertical location predicted neural organization between 70ms and 615ms, and the fragments' category predicted their neural organization between 90ms and 800ms (same as Fig. 2b). b) In a group of central electrodes (C3, TP9, CP5, CP1, TP10, CP6, CP2, Cz, C4, C1, C5, TP7, CP3, CPz, CP4, TP8, C6, C2, T7, T8), vertical location predicted neural representations between 75ms and 595ms, and category predicted the neural organization between 190ms and 585ms. c) In a group of anterior electrodes (F3, F7, FT9, FC5, FC1, FT10, FC6, FC2, F4, F8, Fp2, AF7, AF3, AFz, F1, F5, FT7, FC3, FCz, FC4, FT8, F6, F2, AF4, AF8, Fpz), vertical location predicted neural representations was considerably weaker, but still emerged between 155ms and 230ms, and between 405ms and 430ms, while category predicted the neural organization between 460ms and 765ms. These results show that, in line with differences in visual processing, vertical location information primarily emerges over posterior cortex. Further, category information, which contains the task-relevant distinction between indoor and outdoor scenes, is represented later in more anterior electrodes, which may reflect a progression from perception to decision making and execution. Significance markers denote $p_{corr}$<0.05. Shaded margins represent standard errors of the mean.

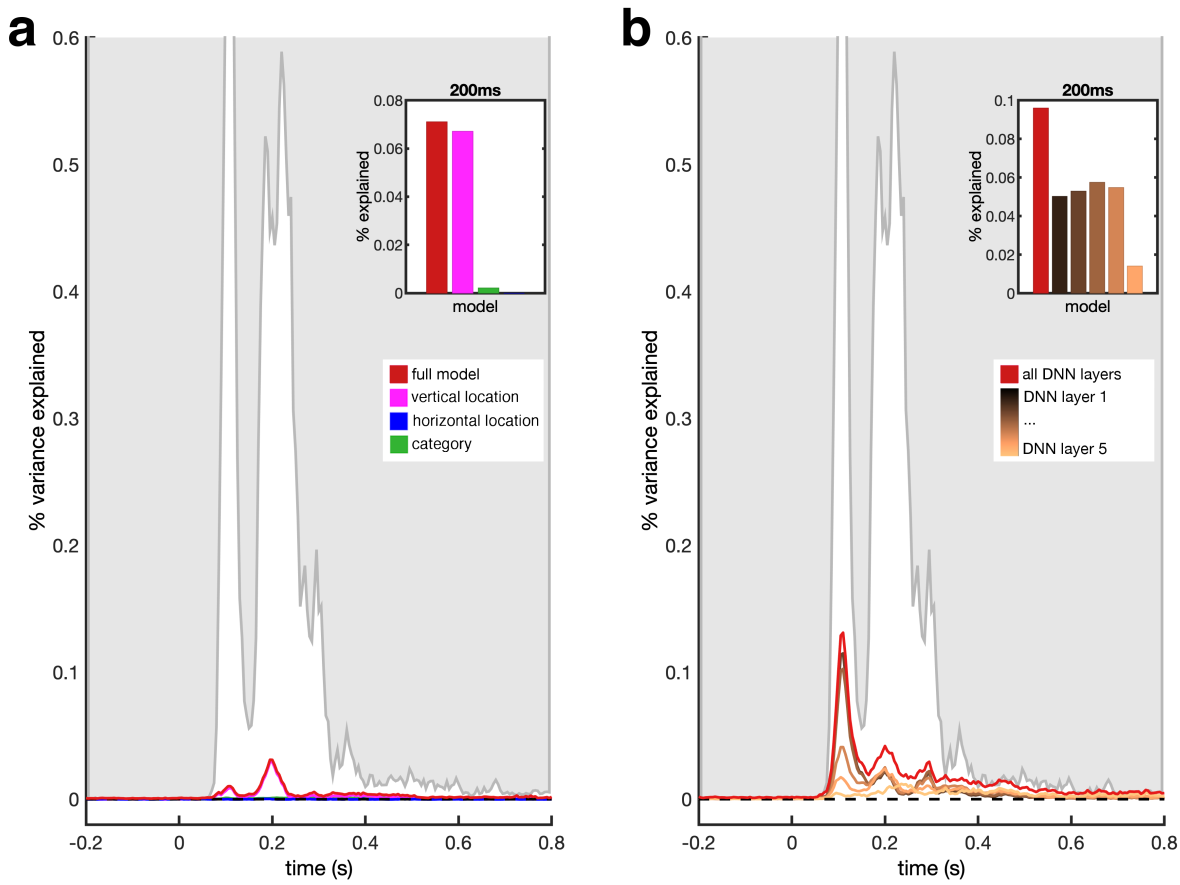


*Figure S2.* Explained variance for different representational models. a) Variance explained ($R^2$) for models containing only the vertical location, horizontal location, or category predictors, and for a model that contains all three predictors. b) Variance explained ($R^2$) for models containing only one of the five DNN layers, and for a model that contains all five layers. The grey line represents an empirical noise ceiling, which provides an estimate of the maximum possible fit given the reliability of the neural data (see Materials and Methods). Inlays show how much of this explainable variance is accounted for by the different models at 200ms post-stimulus, where the greatest vertical-location effect was obtained. Note that the amount of variance explained and the amount of explicable variance are generally low here. This is an expected consequence of our design, which trades a huge stimulus set including a variety of underlying organizational dimensions for the cost of only few (two) repetitions per individual image, thereby yielding noisy estimates for individual RDM entries.

*Table S1.* Peak timings for vertical location information in the model comparison analyses. Intervals reflect the 95% confidence interval (*ci*), which was computed using a bootstrapping method (see Materials and Methods).

| DNN layers included | 1st Peak | 2nd Peak |
|---|---|---|
| **no DNN layers** | 95ms<br>*ci* = [85ms, 120ms] | 200ms<br>*ci* = [197.5ms, 230ms] |
| **DNN layer 1** | 110ms<br>*ci* = [85ms, 115ms] | 195ms<br>*ci* = [190ms, 215ms] |
| **DNN layer 2** | 110ms<br>*ci* = [85ms, 115ms] | 195ms<br>*ci* = [190ms, 210ms] |
| **DNN layer 3** | 110ms<br>*ci* = [85ms, 115ms] | 195ms<br>*ci* = [190ms, 210ms] |
| **DNN layer 4** | 110ms<br>*ci* = [85ms, 110ms] | 195ms<br>*ci* = [190ms, 215ms] |
| **DNN layer 5** | 110ms<br>*ci* = [85ms, 115ms] | 195ms<br>*ci* = [190ms, 215ms] |
| **all DNN layers** | 110ms<br>*ci* = [85ms, 115ms] | 200ms<br>*ci* = [190ms, 215ms] |
